# O-antigen biosynthesis mediates evolutionary trade-offs within a simple community

**DOI:** 10.1101/2023.11.15.566321

**Authors:** Tara C.J. Spencer-Drakes, Angel Sarabia, Gary Heussler, Emily C. Pierce, Manon Morin, Steven Villareal, Rachel J. Dutton

**Affiliations:** Division of Biological Sciences, University of California San Diego, La Jolla, California, United States; Arcadia Science, 3100 San Pablo Avenue, Suite #120, Berkeley, California, United States (current)

## Abstract

Diverse populations of bacteriophages infect and co-evolve with their bacterial hosts. Although host recognition and infection occurs within microbiomes, the molecular mechanisms underlying host-phage interactions within a community context remain poorly studied. The biofilms (rinds) of aged cheeses contain taxonomically diverse microbial communities that follow reproducible growth patterns and can be manipulated under laboratory conditions. In this study, we use cheese as a model for studying phage-microbe interactions by identifying and characterising a tractable host-phage pair co-occurring within a model Brie community. We isolated novel bacteriophage TS33 that kills *Hafnia* sp. JB232 (hereafter *Hafnia*), a member of the model community. TS33 is easily propagated in the lab and naturally co-occurs in the cheese with the Brie community, rendering it a prime candidate for the study of host-phage interactions. We performed growth assays of the *Hafnia*, TS33 and the fungal community members, *Geotrichum candidum* and *Penicillium camemberti*. Employing Random Barcode Transposon Sequencing (RB-TnSeq) experiments, we identified candidate host factors that contribute to TS33 infectivity, many of which are critical to the integrity of bacterial O-antigen. Notably, disruption of these genes results in decreased susceptibility to infection by phage TS33, while simultaneously exhibiting a significant negative effect on the fitness of *Hafnia* in the presence of the fungi. Therefore, O-antigen mutations may have pleiotropic effects on the interactions between *Hafnia* and the rest of the Brie community. Ongoing and future studies aim to unearth the molecular mechanisms by which the O-antigen of *Hafnia* mediates its interactions with its viral and fungal partners.

## Introduction

Bacteriophages are the most abundant replicating entities on the planet. Despite their small size, they possess unique life cycles which provide myriad opportunities to interact with and profoundly impact their hosts. Bacteriophages and their hosts boast an extraordinary genetic diversity, suggesting that host-phage interactions are multi-faceted [1, 2]. The detailed mechanisms of several host-phage interactions have been extensively documented, revealing their robust impacts on host biology. The ecological interactions between bacteriophages and their bacterial hosts occur within the context of a microbial community (microbiome), where phages have significant impacts on host abundance, gene expression, diversification and evolvability [3, 4, 5, 6]. In fact, bacterial processes such as quorum sensing, biofilm formation and metabolism are affected by host-phage interactions [7, 8, 9].

Despite our extensive knowledge on the molecular outcomes of host-phage interactions, current understanding of the impact of community context on these interactions is quite sparse. Moreover, some of the current literature available on the impacts of ecological contexts on host-phage interactions is contradicting [10]. Some studies show that the community may constrain host-phage interactions [11], while others show that the effect of community presence on these dynamics is positive [12, 13]. Similarly, there are no generalised conclusions drawn about the role of community context on host-phage coevolution. In some ecosystems, host adaptation to phage predation has been shown to be inhibited by the community [11, 12] while others have negligible impacts on host adaptation [14] or positive impacts (15, 16).

Notably, time is a limiting factor in these studies as many of these experiments take place over 15 generations on average, which may provide insufficient time to observe ecologically relevant effects [10]. What is more, the microbes under investigation were cultured in an environment that is quite different from their natural habitat. Growth conditions that vary from those in which the microbe has evolved can directly and profoundly impact bacterial and phage growth patterns, thereby potentially producing misleading results. By designing experiments that occur over a realistic ecological timeline, and which closely mimic the fundamental niche of the microbes, we can paint a clearer picture of the rich biology that takes place within microbiomes.

In this study, we employ a model community of cheese microbes to investigate the interactions between phages, their hosts, and the wider microbial community. Cheese surface biofilms, or rinds, are taxonomically diverse microbial communities that follow reproducible growth patterns [17]. Moreover, cheese-associated microbes are easily manipulable in the laboratory. We have previously used a simple Brie microbial community containing the bacterium *Hafnia* sp. JB232, the yeast *Geotrichum candidum,* and the mould *Penicillium camemberti*, to investigate how bacterial-bacterial and bacterial-fungal interactions occur within a community [18, 19]. In this study, we leverage the experimental tractability of this system to study the tripartite interactions between bacteriophages, bacteria, and fungi.

Recently, RB-TnSeq has been applied to the description of host factors required for phage infection. Adler et al. outline a study in *Salmonella enterica* serovar Typhimurium where a dense library was generated and subjected to phage predation [20]. They successfully identified over 300 genes required for infection by multiple phages. This led to the identification of cross-resistance conferred by the RpoS and RpoN genes. Similarly, the genetic determinants of phage resistance were also identified in *E. coli* using the same genome-scale approach [21]. Among the genes required for phage infection were those associated with synthesis of the Gram-negative lipopolysaccharide (LPS) and nutrient transporters. Additionally, a similar method called INSeq was used by Kortight et al. to verify candidate receptors for various coliphages, among which LPS and transport genes were identified [22].

Our primary approach involves using a high-throughput genetic screen that employs barcoded transposon mutant libraries (RB-TnSeq) to (1) discover novel microbial interactions within a community context, and (2) investigate how increasing community complexity affects these interactions. TnSeq methods were originally developed to determine the genetic basis of specific phenotypes. Comparisons of gene frequency under various conditions allows us to highlight key genes that promote and hinder growth [23]. Our lab has uniquely applied these methods to the identification of genetic requirements of bacterial hosts in the midst of a community. By subjecting mutant libraries to various community contexts, we have identified genes, and their associated biochemical pathways, that are important for the growth of *Escherichia coli* or *Pseudomonas psychrophila*. These experiments enabled us to infer interaction mechanisms between *E. coli* or *P. psychrophila* with other community members [18]. Among the important interactions are competition for iron and cross-feeding of amino acids from fungal community members. We also used this system to demonstrate the reorganisation of microbial interactions with increasing community complexity [24].

For the present study, we expand our model Brie community by introducing a novel bacteriophage TS33 isolated from the same type of cheese as the original model community isolates. In effect, we have developed a model host-phage interaction within the already existing model community which has allowed us to study how host-phage interactions occur within a community context. Using a newly-generated barcoded transposon insertion library in *Hafnia* sp. JB232, we investigate the effects of a community of phage TS33 and fungal partners, *P. camemberti* and *G. candidum*, on the growth of *Hafnia*, and vice versa. Over the course of the experiment, we observe that the fungi exert small negative selective pressure on *Hafnia*. However, we witness fluctuations in the fitness of *Hafnia* in the presence of phage TS33 only. Such growth patterns are expected as both the host and phage innovate to preserve their resistance and infectivity, respectively [25]. By the end of the experiment, the abundance of *Hafnia* growing with fungi or phage is very similar to that of *Hafnia* growing alone.

What is striking, however, is the stark decrease in *Hafnia* fitness when both the fungi and the phage are present. Additionally, DNA sequencing reveals that “unfit” *Hafnia* cells are dominated by mutations in genes required for the synthesis of the cell envelope, especially the layer of lipopolysaccharides (LPS). We identify the LPS, and in particular, the O-Specific Polysaccharide (O-antigen) as a key genetic determinant of phage resistance in *Hafnia*. Conversely, mutations in the genes important for O-antigen synthesis offer a negative fitness effect in the presence of the fungi, meaning that these genes are important for resisting negative interactions with fungi. Together, these data highlight the role of the O-antigen in mediating interactions between *Hafnia*, fungi, and phage. Our experiments highlight the importance of community context in the fitness profile of *Hafnia* cells that are interacting with phage TS33, as the outcome of these interactions varies when the fungal partners are included.

In summary, this report illustrates the impact of the ecological context on the molecular outcomes of interactions between bacteria and their phages, and the role of bacterial cell structures in mediating these interactions. The LPS, and specifically, the O-antigen is revealed as a critical interface for bacteria to interact with both phage and fungi within the community. Moreover, by establishing a model host-phage interaction within a community context, we have provided a starting point for full comprehension of the dynamic relationships that exist among fungi, phage, and bacteria. The work presented here is therefore among the first that exposes the molecular underpinnings of host-virus interactions within a community, and the evolutionary trade-offs that exist within a community context.

## Results

### Isolating a lytic bacteriophage that kills *Hafnia* sp. JB232

To establish a model system for studying host-phage interactions within our Brie community, we first isolated a bacteriophage that infects *Hafnia* sp. JB232 from the natural cheese environment of the bacterium. Bacteriophage TS33 was isolated in 2019 from the rind from a different batch of the same cheese that *Hafnia* sp. JB232 was first isolated in 2011 (Figure S1A, [17]). Plaque assays revealed that phage TS33 infects *Hafnia* sp. JB232, and produces a titer of more than 10^10^ pfu/mL following overnight incubation with the host (Figure 1A). The phage TS33 genome was sequenced using Illumina technologies and subsequently analysed using the Viral Proteomic Tree (VipTree) server to determine genome-wide similarities to a bacteriophage reference database [26]. These whole genome comparisons predict that phage TS33 is a member of the viral family *Siphoviridae*. Phage TS33 is most closely related to *Salmonella* phage FSL SP-031 and *Enterobacter* phage phiEap-2, the latter of which infects multidrug-resistant *Enterobacter aerogenes* (Figure 1B).

**Figure 1.**
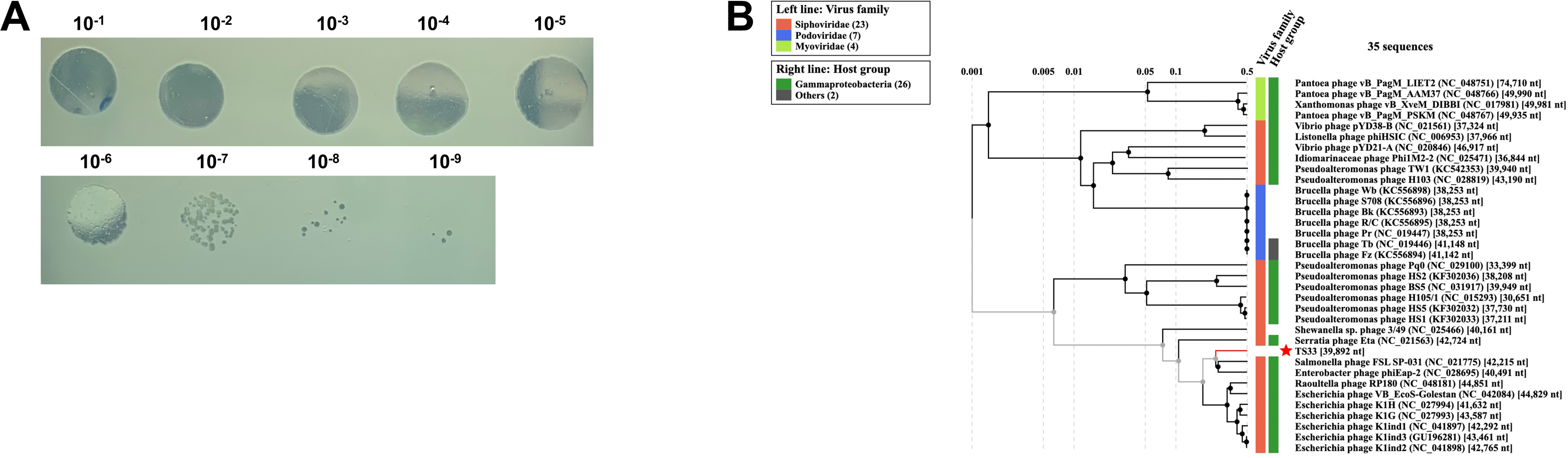
Novel lytic JB232-infecting bacteriophage TS33 is phylogenetically related to members of virus family *Siphoviridae*. (A) Actual plaquing of JB232 phage on *Hafnia* lawn. 3 μL of serially diluted suspensions of pure phage lysates were spotted onto a JB232 lawn. (B) Proteomic tree showing the predicted phylogenetic classification of TS33 among *Siphoviridae*. Phylogeny was determined based on whole genome comparison of the TS33 proteome to the proteomes of well-characterized bacteriophages using the Viral Proteomic Tree (ViPTree) Server.

### Identifying genetic requirements of *Hafnia* growing under different conditions

We used a high-throughput genetic approach to determine how different ecological contexts affect the genetic requirements of *Hafnia* sp. JB232. An RB-TnSeq library was developed in *Hafnia* sp. JB232 by mating with the *Escherichia coli* Keio_ML9 RB-TnSeq library from Wetmore et al. 2015 [27]. The *Hafnia* library contains 103,169 insertion mutations (and thus mutants) in 58,869 distinct locations. Around 88% of the protein-coding genes are disrupted, when considering insertions being in only 10-90% region of those genes (Figure S1B, [28]). Moreover, for each gene represented in the library, there were on average 25.6 different insertion mutants generated (based on the insertion location within the gene).

To determine the effect of ecological context on the genes required for *Hafnia* growth, the pooled transposon mutant library was inoculated on an *in vitro* cheese medium either alone or in the presence of the fungal partners from the Brie community, *Geotrichum candidum* and *Penicillium camemberti*, in a 1:1 ratio (Figure 2A-B). Additionally, each experimental condition was performed with or without phage TS33 (MOI=0.001). On each day of the experiment, cells, spores and phage particles were harvested, diluted, and plated for counting (Figure 2). It is worth noting that under these conditions, it is impossible to assess *Penicillium* growth due to its inability to grow well planktonically.

**Figure 2.**
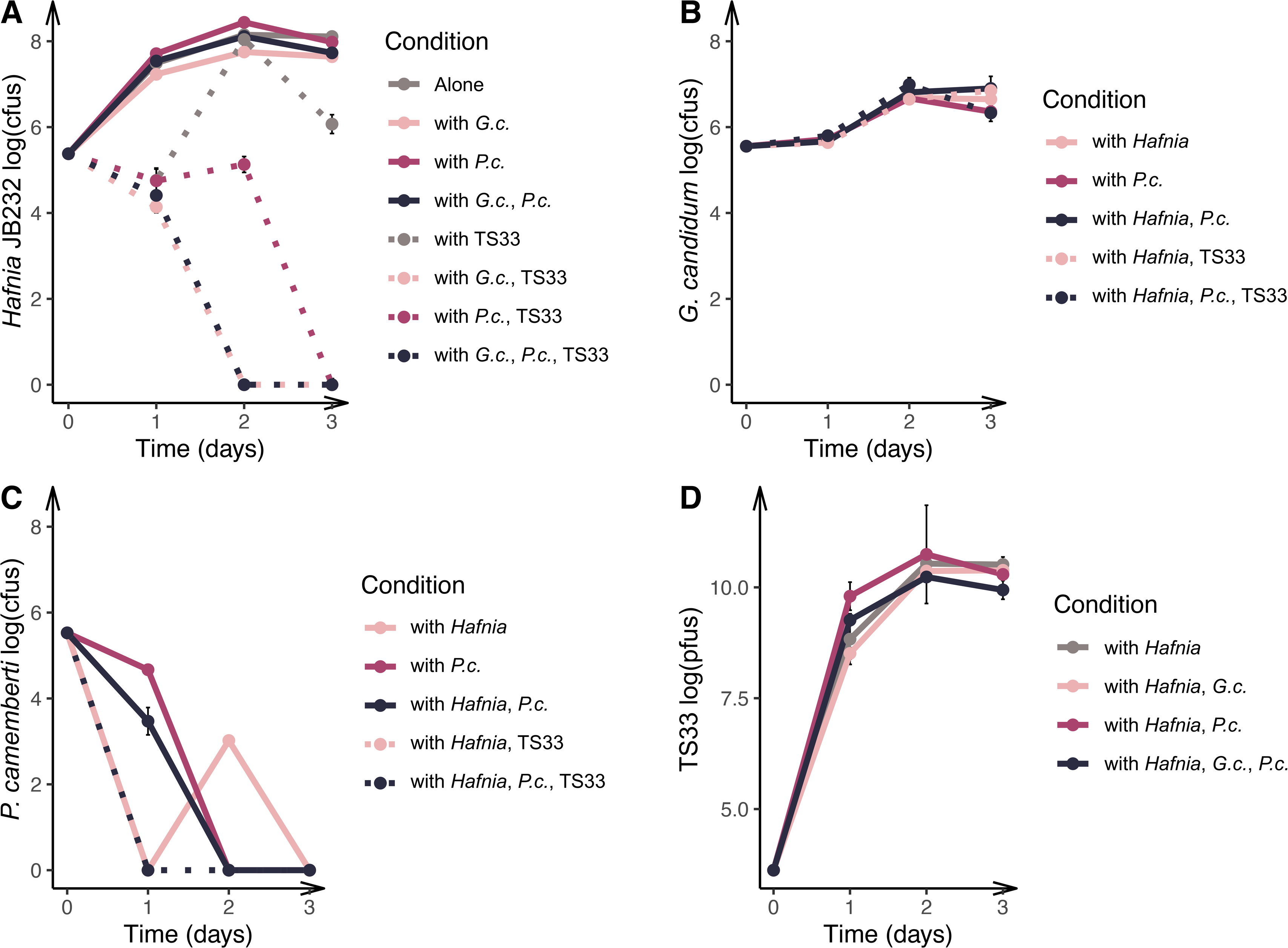
*Hafnia* growth is inhibited in the presence of a community of fungi and bacteriophages. The *Hafnia* sp. JB232 library was grown alone, and with phage and/or fungi on *in vitro* cheese medium for three days. The harvested cells (A), spores (B,C) and phage particles (D) were plated on each day and quantifies using the total number of cfus and pfus respectively.

*Geotrichum* grows steadily over 3 days, attaining levels that are minimally impacted by the presence of bacteria and phage, bacteria only or *P. camemberti* (Figure 2B). Similarly, *Hafnia* and phage TS33 growth levels do not appear to be vastly impacted by the presence of either fungus, or both fungi (Figure 2A). However, in the presence of the phage, the number of *Hafnia* colony-forming units (cfus) decrease on day 1, recover on day 2, and decline again on day 3. Moreover, *Hafnia* growth conditions that include phage TS33 and any combination of the fungi exhibit complete extinction of viable *Hafnia* cells from the community by day 3 (Figure 2A). These data suggest that when both phage and fungi are present, *Hafnia* fails to recover its population levels. Moreover, our data show that phage TS33 density is not amenable to most conditions, except when phage TS33 is interacting with *Hafnia* in the presence of both *G. candidum* and *P. camemberti*. We observe that the community presence significantly decreases the amount of phage particles (Tukey test, p<0.05) after three days of growth. The possibility exists that the extinction of host cells resulted in a decrease in the viability of TS33 virions. However this remains puzzling since the decrease in TS33 over the 3 days is far less than what we observe in the host cells. Nevertheless, our results indicate that the presence of the community negatively affects the growth of both *Hafnia* and phage TS33 (Figure 2A).

To determine the fitness of the different *Hafnia* mutants, we compared the abundance of each mutant in each growth condition after 3 days of growth to its abundance at the start of the experiment. DNA was extracted from the cells in the inoculum and cells harvested on Day 3. RB-TnSeq barcodes therein were amplified using PCR and sequenced. Using sequence data, we calculated the abundances of each barcode and therefore of each insertion mutant at both timepoints using the standard fitness pipeline [27]. For each condition, a fitness value was assigned to each insertion mutant by calculating the log2 change in barcode abundance over 3 days (Figure 3A, Figure S2). Next, we assigned a raw fitness value to each gene represented in the library by calculating the weighted average of the fitness values assigned to all insertion mutants in that gene. The raw fitness for each gene was then normalised in two steps: (1) first, by chromosomal location to account for variance in copy number and (2) second, by assigning neutral fitness of 0 to most genes (for more details, see Materials and Methods and [27]).

**Figure 3.**
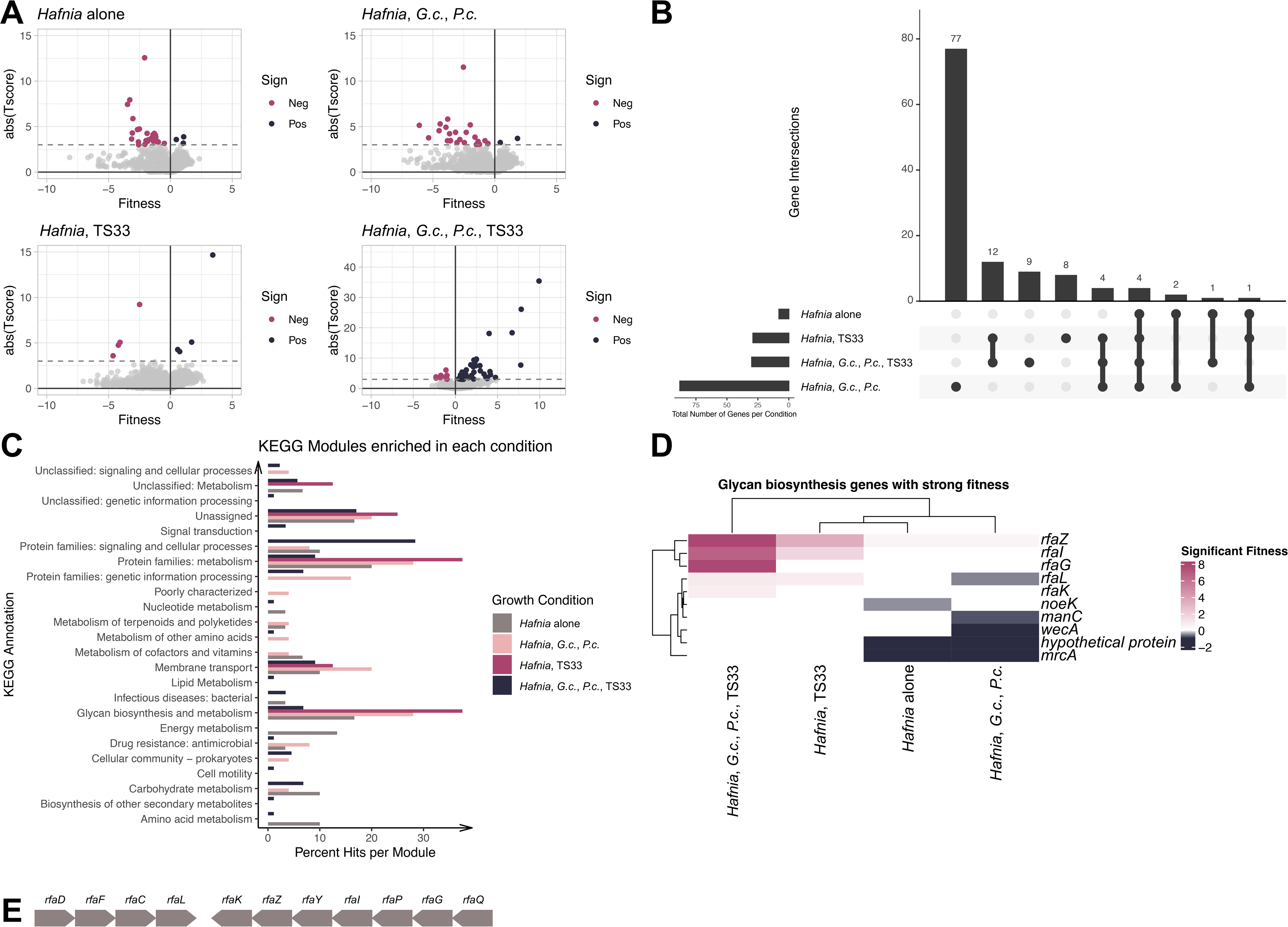
Mutations in genes controlling O-antigen synthesis and ligation may exhibit pleiotropic effects. (A) The barcodes of the *Hafnia* library plated under various growth conditions were amplified and sequenced. The original RB-TnSeq bioinformatic pipeline developed by Wetmore et al. 2015 was used to quantify each barcode and determine the abundance of each mutant in each gene represented in the library. The sum of the abundances of each mutant for each gene was used to assign a fitness value to each gene. Mutations with a significant negative and positive fitness effect are represented in burgundy and blue respectively. (B) The number of genes unique to and shared by the 4 growth conditions were determined using the UpSet R function developed by Lex et al. 2014 (updated by Conway et al. 2017). (C) The KEGG Mapper was used to identify gene pathways enriched under each condition. (D) ComplexHeatMap in R was used to compare the enriched genes under each condition (Gu et al. 2016, Gu 2022). (E) Schematic representation of *rfa* operon in *Hafnia* JB232 Wildtype. Adapted from Pagnout et al. 2019.

Our analysis only included genes whose fitness values were significantly different from 0, as these genes exhibit a significant effect on strain fitness. To this end, a moderated t-statistic (t-score) was calculated for each gene. In every condition, we assumed that most genes with an absolute t-score >=3 had a fitness value that was reliably different from 0 [18]. Thus, our analysis excluded genes that did not meet this criterion. Fitness effects that are greater than 0 represent positive fitness effects; the abundance of these genes at T=3 exceeds their abundance at T=0, as well as the abundance of all other mutants in the library under the growth condition. Conversely, the mutants who decrease in abundance through the experiment experience a negative fitness effect.

Of the 4,021 protein-coding genes represented in the library, we successfully obtained fitness values and moderate t-statistics for 3,863 (96.07%). We employed t-score statistics to filter out genes that lacked a strong fitness effect. In our dataset, this was approximately 96% of all genes (3713/3863). There were 150 genes with strong fitness effects in at least one condition, and 4 of these were conserved across all 4 conditions (Figure 3B, [29], [30]).

Next, we compared the gene functions assigned to these 150 genes among the different ecological contexts of *Hafnia* growth in order to identify whether certain molecular pathways are overrepresented therein. To this end, each gene with a strong fitness effect was annotated using the KEGG Orthology and Links Annotation (KOALA) BLAST and mapped to the KEGG BRITE database (Figure 3C [31]). For each condition, we calculated the percent of the total annotations represented by each KEGG annotation, and then looked specifically for groups that were abundant in all 4 conditions of growth. Genes classified under “Protein families: metabolism” and “Glycan biosynthesis and metabolism” were present in all 4 conditions. It is important to state that 52.9% of the genes classified under “Protein families: metabolism” received the annotation “Glycan biosynthesis and metabolism” or the sub-annotation “Lipopolysaccharide biosynthesis proteins”. Moreover, the glycan biosynthesis and metabolism genes are all involved in lipopolysaccharide (LPS) biosynthesis. The LPS is a glycolipid that is comprised of three parts - lipid A, the core polysaccharide, and the O-antigen which is the most outward-facing part of the structure. The LPS is well-documented as a critical cell structure for combatting attacks from bacteriophages, other bacteria, and chemicals [32, 33]. Specifically, it is a common adsorption site for diverse bacteriophages [34, 35]. Here, we see that the LPS genes are present in some way under all conditions of growth.

To determine the specific impacts of these genes in each condition, we investigated the fitness effects of each gene associated with glycan/LPS biosynthesis and metabolism. The LPS biosynthesis genes were primarily annotated as *rfa* genes (known players in LPS biosynthesis in *E. coli*, [36]) and glycosyltransferases predicted to be involved in cell wall biosynthesis (Figure 3E, adapted from [37]).

Insertions in the *rfaL* gene, which synthesises the O-antigen ligase, exhibit positive fitness effects in all conditions containing phage, and negative fitness effects in the presence of fungi only (Figure 3A, Figure 3D, [38], [39]). These fitness effects were stronger than those observed when *Hafnia* grows alone. Interestingly, disruption of the *rfaZ* (KDO-transferase) gene has a relatively small positive effect on the fitness of *Hafnia* growing with fungi; however, this effect is three times greater when *Hafnia* grows with phage TS33 and is seven times greater when *Hafnia* grows in the community (Figure 3A, Figure 3D). Another operon encoding predicted glycosyltransferases that contribute to cell wall biosynthesis follows a similar pattern; overall, we observe that mutations in any genes from this operon have negative fitness effects on *Hafnia* growing with fungi, and positive fitness effects during growth with phage or the whole community. In other words, in the presence of TS33, disrupting LPS biosynthesis appears to provide a fitness benefit to *Hafnia* but may offer a fitness cost in the presence of the fungi.

It is worth noting that the positive fitness effects that we observe in these genes when *Hafnia* is growing with the community may be attributed to strong selection for them by the phage. Subsequently, they succumb to potentially negative interactions with the fungi, evidenced by the stark decrease in the growth of *Hafnia* in the community context compared to all other conditions. Thus, we hypothesised that some genes involved in LPS synthesis might have pleiotropic effects in *Hafnia*.

### Determining the role of LPS in host-virus-fungus interactions

To test our hypothesis, we used a high titer of phage to select for phage-resistant *Hafnia* mutants (Figure 4A). We successfully isolated mutants only in genes involved in LPS and specifically, O-antigen biosynthesis: *rfaL, rfaH, manC* (mannose-1-phosphate guanylyltransferase), and *wzy* (a predicted O-antigen polymerase gene). RfaH inhibits the termination of transcription of LPS biosynthesis genes [40], while ManC (CpsB in *E.coli*) is involved in the synthesis of the growing O-antigen.

**Figure 4.**
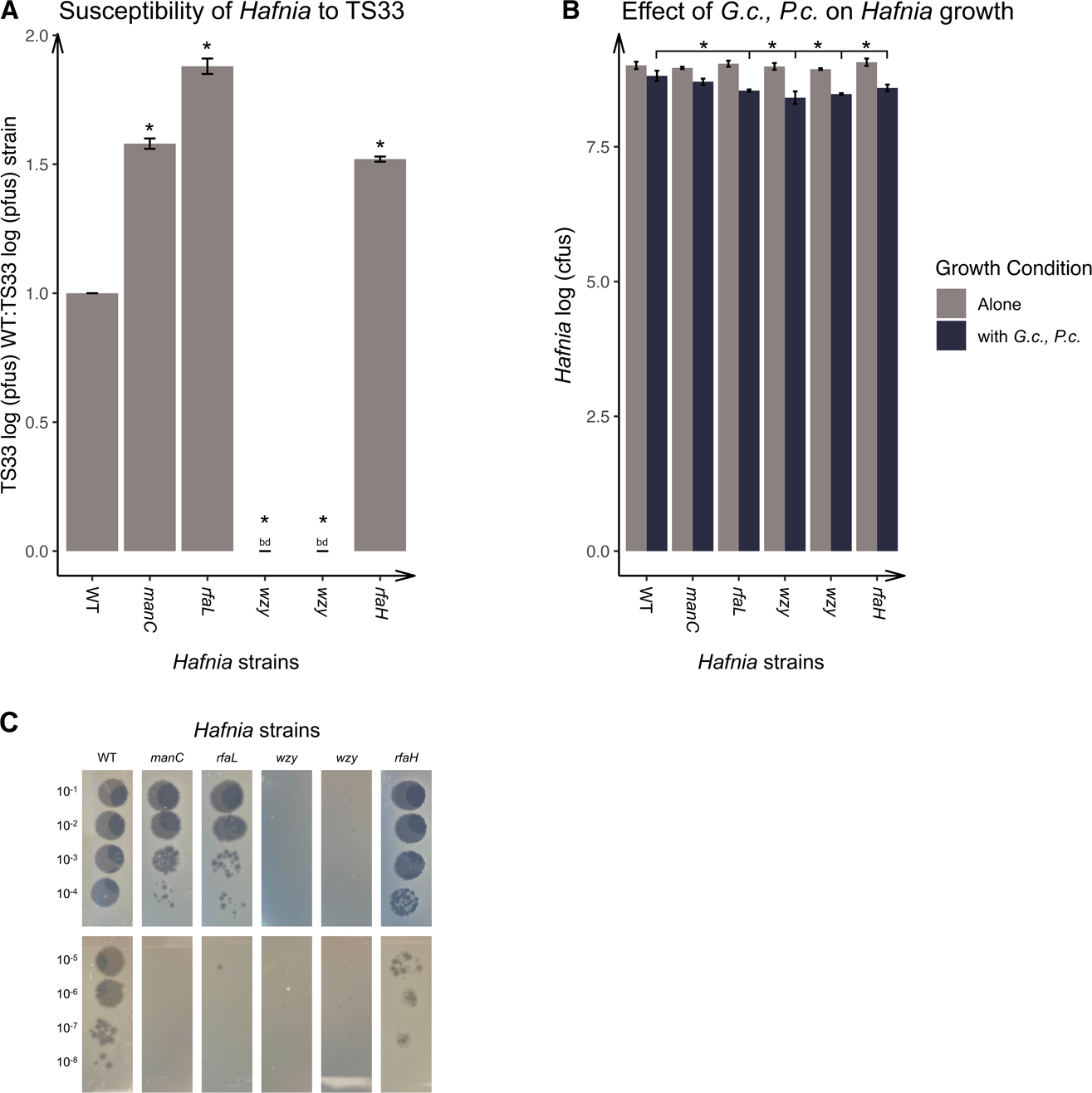
LPS mutations increase JB232 resistance to TS33 infection but decrease bacterial growth in the presence of the fungi. (A) Comparing phage infectivity of WT and mutant strains. Phage infectivity is measured by the number of TS33 phage particles (shown here as the number of plaque forming units/pfus) produced following infection on a soft agar lawn of each bacterial strain. (B) Effects of fungi on growth of *Hafnia* strains over 3 days, measured in the number of viable bacterial cells (represented here in colony forming units (cfus). (C) Images of phage plaquing on various *Hafnia* mutants.

We hypothesised that the susceptible insertion mutants within the *Hafnia* library growing with phage TS33 experience high levels of phage predation, resulting in the community being dominated by TS33-resistant mutants and accounting for the spike in *Hafnia* counts on Day 2. However, eventually, the *Hafnia* counts recover, possibly due to the emergence of counter-resistant mutations in phage TS33. However, in the presence of either fungus or both fungi, the phage TS33-resistant mutants experience a negative fitness, resulting in their decrease in growth in any ecological context containing fungi (Figure 2A, [25]).

To directly test this hypothesis, we aimed to determine whether phage TS33-resistant mutants with disrupted LPS genes experience a fitness deficit in the presence of both fungi. We subjected the *Hafnia* library to high titers of phage TS33 for 24 hours in liquid LB. The remaining viable cells were plated on solid LB media and the emergent colonies were picked and struck three times to purify the strains. Phage TS33 was diluted and titered on top agar lawns of each mutant. In total, we isolated and sequenced the barcodes of 50 mutants, each displaying decreased or no susceptibility to the TS33 lysate compared to the wild-type (Supplementary Table 1). Sequencing of the mutant barcodes revealed disruptions in the genes associated with biosynthesis of the cell wall, especially of the LPS. From the 50 mutants, we selected 5 that were represented in our RB-TnSeq results in at least one of the 3 experimental conditions: one in *manC*, one in *rfaL*, one in *rfaH*, which transcriptionally controls the expression of other *rfa* (LPS-synthesising) genes, and two in *wzy*, which encodes an O-antigen polymerase (Figure 4A, 41). The mutant and wild-type *Hafnia* cells were then inoculated on *in vitro* cheese medium with or without the fungal community members. After 3 days, the cells were harvested and quantified. For each *Hafnia* strain plated, we compared the number of cfus after growth with fungi to that after growth alone (Figure 4B, Figure S3A).

The *wzy* mutants were completely resistant to infection by phage TS33, while the *manC*, *rfaL*, and *rfaH* mutants exhibited drastic reductions in their susceptibility. From these results, we can observe a fitness benefit to the *Hafnia* cells by the disruption of genes involved in O-antigen synthesis, with the strongest benefit coming from the interruption of the O-antigen polymerase gene (Figure 4A, Figure S3B). Interestingly, in the presence of the fungi, the *rfaL, rfaH* and *wzy* mutants experience a significant decrease in growth, compared to the wild-type (Figure 4B, Tukey test, p <0.05), strongly suggesting a role played by the O-antigen in the modulation of interactions between *Hafnia* and its viral and fungal environment. This fitness defect is not observed in the *manC* mutant, although *manC* is also involved in O-antigen synthesis [42]. The failure of the disruption of *manC* to negatively affect *Hafnia* fitness in the presence of the fungi suggests that these specific sugar residues are not required for *Hafnia*-fungi interactions.

## Discussion

Most studies of bacteria-phage interactions completely exclude the natural community wherein the microbes exist naturally. In this study, we used *Hafnia* sp. JB232 interactions with phage TS33 as a model to uncover the principles of community impact on host-phage interactions. Under each condition of growth, we quantified the number of bacterial cells to understand the effects of different combinations of community members on *Hafnia* growth. Moreover, using a high-throughput genetic screen, we were able to further investigate these growth effects by studying the fitness landscape of the *Hafnia* cells in each growth condition. Our analyses reveal that phage can lead to fluctuations in *Hafnia* growth that eventually the bacteria recover from. During the time period of our experiment, neither the phage nor fungi alone elicited as drastic a depletion in the final counts of *Hafnia*, as both of them did together. In other words, neither group of community partners is sufficient to produce a strong negative outcome on *Hafnia* growth; both are required for extinction of *Hafnia*.

Similar and contradicting observations have been made by other scientists. In a study of wastewater microbial ecology, Johnke et al. unearthed suppressive effects of a microbial community on *Klebsiella* sp. [11]. Under phage selective pressures, the bacterium experiences more severe impacts on its growth. In the same study, they also show that the presence of some other specific community members contributes to the sustained growth of *Klebsiella*. Conversely, *P. fluorescens* and *P. aeruginosa* both benefitted from a community presence amidst phage predation [12, 13]. While our results most closely support those projected by Johnke et al., it is probable that the community effects are dependent on community composition. Our studies involved the use of fungi, *P. camemberti* and *G. candidum*, which are known to have negative interactions with other microbes. *P. camemberti* produces effects on *E. coli* that are similar to those exhibited by a beta-lactam antibiotic [19]. Moreover, *G. candidum* is believed to exert oxidative stress on *E. coli* (18). In this previous study, in the presence of the fungus, *E. coli* depends heavily on genes such as *acrA* and *acrB* that enable it to resist oxidative stress. Moreover, in the presence of the community, these genetic requirements are alleviated, suggesting that activity from one of the community members is either preventing oxidative stress or empowering *E. coli* to deal with it. It is possible that these molecular products of both fungi produce a negative effect on *Hafnia* mutants resistant to phage infection. To further understand the relationship between community presence and growth of focal bacteria, it is essential to investigate the molecular mechanisms that underlie their interactions.

Our study revealed a unique role of the LPS, specifically the O-antigen, in *Hafnia*’s interactions with its community. Specifically, mutations in genes involved in O-antigen synthesis (*rfaL*, *wzy*) and regulation of the LPS biosynthesis gene expression (*rfaH*) produced a growth deficit in the presence of fungi and increased growth in the presence of phage, compared to the wild type. The LPS is a major component of the defence mechanism of Gram-negative bacteria, as it protects them from chemical attacks by other microbes and the environment. The intact LPS is often exploited by phage during attachment to the host [34]. In fact, the O-antigen has been shown to be a specific target of several bacteriophages [43]. Additionally, phages and antimicrobials are known to negatively affect bacterial cells with compromised LPS [44, 45]. Burmeister et al. demonstrate that LPS mutations confer phage resistance but restore antibiotic susceptibility [45]. We observe that *Hafnia* LPS mutants develop resistance to phage TS33 infectivity while experiencing growth deficits in the presence of the fungi. In light of the negative interactions that these fungi are known to have with other microbes, it is possible that the antibiotic/oxidative stress that they apply accounts for the growth defect in LPS mutants growing under these conditions. In other words, genes controlling LPS biosynthesis may have pleiotropic effects in *Hafnia,* and lead to trade-offs with evolutionary consequences for *Hafnia* growing in a community of fungi and phage. Moreover, genes involved in the synthesis of Enterobacterial Common Antigen (ECA) have been shown to be important in interactions within increasingly complex microbial communities [24]. Within the context of our data, these findings suggest a general role of glycan-modified antigens in community interactions.

There is additional evidence that the negative effects of antibiotics on LPS-deficient cells may be genotype-specific. In a study by McGee et al., 3 of 4 LPS mutants under study were resistant to both phage and antibiotics of various classes [46]. They observe that *rfaH* and *yciM* (regulating synthesis of Lipid A; Sunayana and Reddy, 2013) mutants are phage and antibiotic-resistant; the phage-antibiotic resistance trade-off was observed only in the *rfaP* (facilitating synthesis of the core oligosaccharide) mutant. While we were not able to isolate an *rfaP* mutant for our experiments, our findings do suggest that all LPS mutations may not provide a trade-off within a community context. *manC* is involved in the biosynthesis of GDP-mannose, a key sugar nucleotide precursor used in the synthesis of the O-antigen [47]. However, the mutation in *manC* was not sufficiently deleterious to produce a statistically significant growth deficit in the presence of fungi. Conversely, insertion mutants in O-antigen polymerase (*wzy*), O-antigen Ligase (*rfaL*) and the anti-termination protein that regulates expression of *rfa* genes (*rfaH*) exhibit growth deficits in the presence of the fungi. We expect that mutations in these genes would result in improper polymerization of the sugar residues within the O-antigen, faulty attachment of the O-antigen to the growing LPS and poor regulation of LPS synthesis. In any case, we would predict that these mutants lacked the ability to assemble and ligate the O-antigen on a whole, whereas the *manC* mutation only affects one residue. Thus, it is plausible that the sugar residues introduced by manC specifically have no bearing on host susceptibility to negative interactions with the fungi. Combining our results with those of McGee et al., it is plausible that some Lipid A and the O-antigen mutations are likely implicated in increased susceptibility to antibiotics and other negative interactions with community members [46]. However, the O-antigen is possibly more important than Lipid A for the existence of phage resistant trade-offs when experiencing the aforementioned negative environmental stimuli.

Furthermore, our unique approach to the study of host-phage interactions underscores the critical importance of considering the community in these investigations. Using *Hafnia* as a model, we show that the ecological context can affect the growth outcome of bacteria. The outcome of *Hafnia*-TS33 interactions is more positive when the fungi are absent. Additionally, the fungi have a negative effect on the fitness of *Hafnia* that is not observed in the growth condition with the fungi. The phage also experienced density suppression mediated by the fungi. These results are similar to what is seen in wastewater microbiomes, where protists negatively impact host-phage interactions. Therefore, including all members of the community in our study allowed us to observe the dependence of *Hafnia*-phage interactions on both fungi.

Altogether, our findings shed light on the unique contributions of community members to bacteria-phage interactions, and thus, reinforce the need to consider the microbiome in studies of the biology of bacteria and phages. Additionally, through this study, we have just begun to understand the role of the LPS in mediating host-phage interactions within a community. Finally, we have introduced a model host-phage interaction within a community context, which we hope will serve as a platform for future investigations of microbial interactions across the three domains of life.

## Materials and Methods

### Media preparation

Community growth assays were performed using 10% cheese curd agar at pH 7 (10% freeze-dried Bayley Hazen Blue cheese curd (Jasper Hill Farm, VT), 3% NaCl, 0.5% xanthan gum, and 1.7% agar); 10M NaOH was used to buffer the pH from 5.5 to 7 [17]. All growth assays and inoculations were performed at room temperature.

### Bacterial, fungal, and phage strain selection and preparation

All bacterial and fungal isolates were obtained from a natural rind cheese as previously described [17]. The community was selected based on its success as a model in a prior study conducted in the group [18].

As *Hafnia* sp. JB232 is the only bacterium in the model community, it was used to isolate phages from a different batch of the natural rind cheese from which it was originally isolated. The cheese rind was scraped and homogenised in SM buffer (100 mM NaCl, 8 mM MgSO_4_, 50 mM Tris-Cl). The suspension was vortexed vigorously and centrifuged at 13000 rpm, at 4°C. The supernatant was filtered using a 0.45-μM filter. The filtrate was then serially diluted; each dilution was mixed with 200 μL of a late log culture of *Hafnia* sp. JB232 and 4.5 mL soft agar (0.05% Bacto-agar, 25% LB agar), and poured onto solid LB media. After 24 hours, the resulting plaques were picked and struck out 3 times on solid LB medium with *Hafnia* sp. JB232. On the third quadrant streak, one plaque was picked and cultured in liquid LB medium with a colony of *Hafnia* sp. JB232. After 16-18 hours of incubation, the cells were pelleted and the supernatant was filtered using a 0.45-μM filter. This phage lysate was stored at −80°C with fresh 1XPBS-40% glycerol.

To prepare stocks of *P. camemberti* and *G. candidum*, spores were obtained from Danisco and resuspended in 1XPBS-Tween0.05%. Spore stocks of equal volume were aliquoted with fresh 1XPBS-40% glycerol and stored at −80°C. To quantify the stocks, aliquots were frozen for 48 hours, after which they were thawed and plated on solid LB medium. For each fungal strain, we calculated the number of cfus per milliliter of strain stock.

### Transposon mutant library construction in *Hafnia* sp. JB232

*Hafnia* sp. JB232 was mutagenized by conjugation with *E. coli* strain APA766 (donor WM3064 which carries the pKMW7 Tn5 vector library containing 20 bp barcodes) [27]. In pilot experiments, improved conjugation efficiencies were observed when the donor and recipient strains were at the mid-log phase prior to conjugation. The APA766 donor strain was grown in LB-kanamycin:diaminopimelic acid (DAP) (1 mL frozen stock into 99 mL of LB with 50 μg/mL kanamycin and 60 μg/mL DAP) at 37 deg C at 200 rpm until the culture reached mid-log phase. *H. alvei* sp. JB232 was grown in LB at 30 deg C at 200 rpm until the culture reached the mid-log phase. *E. coli* donor cells were pelleted and washed twice with 100 mL of LB without antibiotics. Donor and recipient cells were mixed at a 1:1 cell ratio based on OD600 measurements, pelleted, and resuspended in 100 μL. 40 μL of the mix was plated on a nitrocellulose filter on an LB plate containing 60 μg/mL DAP. Conjugation took place at 30 deg C overnight. 8 conjugations were performed. The conjugations were each resuspended in 2 mL of LB broth and then 100 ul was plated on an LB:kanamycin (50 μg/mL) agar plate. This was done for a total of 120 selection plates. Plates were left at room temperature until single colonies formed (about 3 days). The resulting transconjugants were scraped into LB from the selection plates and pooled together. The resulting pool was mixed well and then diluted back to an OD600 of 0.2 in liquid LB:kanamycin (50 μg/mL). The pool was then grown to mid-log phase, and two 5 mL samples were taken and pelleted. Cell pellets were frozen at −80 deg C for later library characterization. Sterile glycerol was then added to a final concentration of 15%. 1-mL glycerol stocks were stored at −80 deg C. The final library was estimated to contain ∼160,000 mutants.

### Library preparation and sequencing

Genomic DNA was extracted from the cell pellet of the *Hafnia* sp. JB232 mutant library using Phenol:Chloroform:Isoamyl alcohol (ph 8) [18]. Library preparation was conducted by the standard ≥100 ng protocol from the NEBNext Ultra II FS DNA Library Prep Kit for Illumina (NEB) with minor modifications. 500 ng of input DNA and a 7-minute fragmentation incubation were used. For adaptor ligation, a custom splinkerette adaptor prepared at 15 μM was used (duplexed DNA of /5Phos/GATCGGAAGAGCttttttttttcaaaaaaaa/GTGACTGGAGTTCAGACGTGTGCTCTTCCG ATC*T). The USER enzyme step was not performed. For size selection, 0.15X (by volume) NEBNext Sample Purification Beads (NEB) were used for the first and second bead selection steps. Before enrichment, DNA was digested with BsrBI (NEB) for 20 minutes at 37 °C prior to heat inactivation at 80 °C for 20 min. The DNA was then purified using 1X AMPure XP beads (Beckman Coulter) in preparation for PCR enrichment. The NEBNext Ultra II FS DNA Library Prep Kit for Illumina (NEB) PCR protocol was used, with custom primers and 30 cycles for the PCR step (P6_ET-Seq2_3_R1: AATGATACGGCGACCACCGAGATCTACACGTCGTCacacTCTTTCCCTACACGACGCTC TTCCGATCTNNNNNNGATGTCCACGAGGTCTC*T and P8_ET-Seq3_R2: CAAGCAGAAGACGGCATACGAGATACATCGGTGACTGGAGTTCAGACGTGT*G). Following library preparation, PE150 sequencing was done by Novogene on the Novaseq 6000 platform. 15 GB of sequencing data was received.

### Library characterization

Sequencing reads were analysed using the Perl script MapTnSeq.pl from Wetmore et al. 2015 [27]. This script maps reads to the *Hafnia* sp. JB232 genome and is publically available at https://bitbucket.org/berkeleylab/feba. The script DesignRandomPool.pl was used to generate the file of mapped barcodes. There were a total of 165,694 insertions mapped, with 103,169 located within the central 10-90% of a gene. There were central insertions in 88% of protein-coding genes.

### *In vitro* community experiment using RB-TnSeq library in *Hafnia* sp. JB232

#### Library pre-culture

The *Hafnia* sp. JB232 transposon library was thawed on ice, and cultured in liquid LB with kanamycin (1 μg/mL) to the mid-log phase. 5 mL of the preculture was pelleted and stored at −80°C to be used as the T0 reference in the fitness analysis.

#### Inoculations

The remaining cells and thawed spores were pelleted and washed in 1XPBS/Tween before inoculation. Respective to the condition of growth, 2.4×10^5^ cfus of *Hafnia*, 3.37×10^5^ cfus *P. camemberti*, 3.59×10^5^ cfus *G. candidum,* and 4.18×10^3^ pfus TS33 were inoculated on CCA plates using sterile glass beads.

#### Harvest

At T=24h, 48h, and 72h, cells, spores, and phage particles were harvested from the fitness assays. For each harvest, CCA plates were flooded with 2-3 mL of 1XPBS-Tween0.05% and cells were gently scraped off. Cells and/or spores were pelleted, and the pellets were washed twice. In conditions containing phage, the supernatants generated after each wash were combined. Before quantification, cells and supernatants from each fitness assay were aliquoted and stored in 1XPBS-40% glycerol at −80°C before quantification. The remainder was stored at −80°C without glycerol for DNA extraction.

#### Growth quantification

Cells/spores in frozen stocks were pelleted by centrifugation and washed with 1XPBS-Tween0.05%. The cells were resuspended in 200 μL 1XPBS-Tween0.05%, serially diluted, and spread evenly on solid LB media using glass beads. Frozen aliquots of phage were thawed, serially diluted, and titered on a soft agar lawn of *Hafnia* (as described above). The number of colonies and plaques were used to calculate the number of colony-forming and plaque-forming units, respectively, in 1 milliliter of harvested material. Subsequently, we calculated the total number of cfus and pfus harvested. These values were analyzed for statistical significance using the Tukey’s test. q-q-plots, the Shapiro-Wilk (S-W) test and Pearson Chi-Squared Normality tests were used to test for normality using R version 4.2.2.

#### gDNA Extraction

From each fitness assay, gDNA was extracted using Phenol:Chloroform:Isoamyl alcohol (ph 8). Cell lysis was conducted by adding a homemade lysis buffer (10 mM Tris-Cl pH 8, 100 mM EDTA pH 8, 1% SDS, 10 μg/mL RNAse A, 1 mg/mL Lysozyme) to the pellet, vortexing the tubes at maximum speed and incubating at 37°C for 1 hour. An equal volume of Phenol:Chloroform:Isoamyl alcohol was added to the lysate, after which the tubes were centrifuged at maximum speed for 15 minutes at 4°C. To precipitate the gDNA, 0.1 volume of 10M ammonium acetate and an equal volume of extremely cold isopropanol was added to the aqueous phase (upper layer). After centrifuging for 3 minutes at maximum speed, the pellet was washed with fresh 70% ethanol and resuspended in 25 µL of DNAse/RNAse-free water overnight.

#### Barcode Amplification and Sequencing

After resuspension, the DNA was quantified via Qubit dsDNA HS assay kit (Invitrogen). Subsequently, we used the BarSeq PCR protocol devised by Wetmore et al. 2015 to amplify only the barcoded region of the transposons [27]. The PCR reaction and program used were also previously reported by Morin et al. 2018 [18]. In total, we performed 13 PCRs (T0 sample and 12 harvest samples) involving 13 different multiplexing indexes. 10 µL of each PCR product was pooled, after which the entire pool was purified using the Qiagen MinElute purification kit and quantified via Qubit dsDNA HS assay kit (Invitrogen) and sequenced at Novogene on 1 HiSeq PE 150 lane (6 bp, i7 single index, Illumina). The output of the lane was 375 million reads.

#### RB-TnSeq data processing and fitness analysis

To calculate fitness values for each gene represented in the library, the de-multiplexed barcode reads were processed using the RB-TnSeq data analysis pipeline assembled by Wetmore et al. 2015 [27]. Each script is available to the public at https://bitbucket.org/berkeleylab/feba. The pipeline quantifies the number of reads associated with each barcode to determine the abundance of each insertion mutant under each condition, at T=0 and T=3. Reads that do not map to the characterised barcodes in the library are filtered out, as well as barcodes that are counted less than 3 times in the T0 sample. Moreover, genes represented by less than 30 barcode reads are excluded from further analysis. For the barcodes that meet these parameters, a raw fitness value (log2 change in abundance over 3 days) is calculated and assigned. The weighted average of the fitness values of all insertion mutants in a single gene is calculated as the raw fitness value in that gene. The raw fitness value of each gene is subsequently normalised in two stages. First, the smoothed median is subtracted from each fitness value. This calculation is essential to account for variance in copy number as genes closer to the replication fork have a higher copy number. Second, we assume that most insertions have neutral fitness effects, or fitness effects of 0. To this end, the mode of the fitness distribution is subtracted from each fitness value (for more details, see Wetmore et al. 2015 [27]).

Because we assume that most genes have neutral fitness, our analysis aims only to highlight genes with strong fitness effects, that is, genes whose fitness values were reliably different from 0. A moderate t-statistic (t-score) was calculated for each gene as the ratio of the gene fitness to its standard deviation (for more details, see Wetmore et al. 2015 [27]). T-scores were further used to filter genes with neutral fitness from the analysis. Specifically, we assumed that most genes with reliable neutral fitness had a t-score less than 3, and thus were excluded.

For the remaining genes, we calculated an average fitness score and t-score for the three replicates in each condition. Genes were denoted as having strong positive fitness effects if their abundance of these genes at the end of the experiment exceeds their abundance in the inoculum, as well as the abundance of all other mutants in the library under the growth condition. These genes also had a fitness value that was greater than 0. On the other hand, negative fitness effects are observed in genes whose insertion mutants exhibit a strong growth deficit after 3 days of growth.

### Selection and isolation of TS33-resistant mutants for fitness assay

TS33-resistant mutants were isolated from the RB-TnSeq library in two ways. First, the library was pre-cultured to the late-log phase in liquid LB medium with kanamycin (1 μg/mL) and 1.5×10^8^ pfus TS33. The cells were pelleted, washed twice, and resuspended in 1XPBS-Tween0.05%. The cells were then diluted and each dilution was plated on solid LB media using sterile glass beads. 8 colonies were picked and struck twice on solid LB media. Titer assays of TS33 on soft agar lawns on these strains showed that they are completely resistant to TS33 infection. Whole genome sequencing was conducted at SeqCenter, LLC. Two *wzy* mutants were selected from this set.

Second, the library was pre-cultured to the late-log phase in liquid LB medium with 1.5×10^8^ pfus TS33 and no kanamycin. The cells were pelleted, washed twice, and resuspended in 1XPBS-Tween0.05%. The cells were then diluted and each dilution was plated on solid LB media using sterile glass beads. 42 colonies were picked and struck twice on solid LB media.

Titer assays of TS33 on soft agar lawns on these strains showed varying levels of resistance across the mutants, with some being completely resistant and others showing significantly decreased susceptibility to TS33 infection. Barcode amplification was conducted as previously described (see *above*). Whole genome sequencing was conducted at SeqCenter, LLC on three mutants in the genes *rfaL, rfaH, manC*.

### *In vitro* fitness assay using insertion mutants in *Hafnia* sp. JB232

#### Inoculation

The 5 selected mutant strains and the WT *Hafnia* sp. JB232 were pre-cultured to late-log phase in liquid LB medium. The cells were pelleted by centrifugation and washed twice with and resuspended in 1XPBS-Tween0.05%. Frozen stocks of *G. candidum* and *P. camemberti* were washed twice with and resuspended in 1XPBS-Tween0.05%. For each strain, ∼1.08×10^8^ cfus on average were plated on CCA medium, with or without both fungi (3.82×10^6^ cfus of *G. candidum* and 9.82×10^5^ cfus *P. camemberti* were plated).

#### Harvest

After 72 hours, the cells and spores were harvested using 1XPBS-Tween0.05% as previously described.

#### Quantification of growth

The harvested cells and spores were diluted in 1XPBS-Tween0.05% and plated on solid LB medium. *Hafnia* colonies were counted and the number of cfus harvested was calculated for each mutant under each condition of growth (alone or with fungi). These values were analysed for statistical significance using the Tukey test in Microsoft Excel, and the Shapiro-Wilk (S-W) and Pearson Chi-Squared tests as well as q-q plots were used to test for normality.

## References

1. Kauffman KM, Chang WK, Brown JM, Hussain FA, Yang J, Polz MF, Kelly L. Resolving the structure of phage–bacteria interactions in the context of natural diversity. Nat Comm. 2022;13:372.

2. Jessup CM, Forde SE. Ecology and evolution in microbial systems: the generation and maintenance of diversity in phage-host interactions. Res Microbiol. 2008;5:382–89.

3. López-Leal G, Camelo-Valera LC, Hurtado-Ramírez JM, Verleyen J, Castillo-Ramírez S, Reyes-Muñoz A. Mining of Thousands of Prokaryotic Genomes Reveals High Abundance of Prophages with a Strictly Narrow Host Range. mSystems. 2022;7:e0032622.

4. Thompson LR, Zeng Q, Chisholm SW. Gene Expression Patterns during Light and Dark Infection of *Prochlorococcus* by Cyanophage. PLoS One. 2016;11:e0165375.

5. Kortright KE, Chan BK, Evans BR, Turner PE. Arms race and fluctuating selection dynamics in *Pseudomonas aeruginosa* bacteria coevolving with phage OMKO1. J Evol Biol. 2022;35:1475–87.

6. Cairns J, Frickel J, Jalasvuori M, Hiltunen T, Becks L. Genomic evolution of bacterial populations under coselection by antibiotics and phage. Mol Ecol. 2017;26:1848–59.

7. Sharon I, Alperovitch A, Rohwer F, Haynes M, Glaser F, Atamna-Ismaeel N, Pinter RY, Partensky F, Koonin EV, Wolf YI, Nelson N, Béjà O. Photosystem I gene cassettes are present in marine virus genomes. Nature. 2009;461:258–262.

8. Shah M, Taylor VL, Bona D, Tsao Y, Stanley SY, Pimentel-Elardo SM, McCallum M, Bondy-Denomy J, Howell PL, Nodwell JR, Davidson AR, Moraes TF, Maxwell KL. A phage-encoded anti-activator inhibits quorum sensing in *Pseudomonas aeruginosa*. Mol Cell. 2021;81:571–583.e6.

9. Fernández L, González S, Campelo AB, Martínez B, Rodríguez A, Pilar García P. Low-level predation by lytic phage phiIPLA-RODI promotes biofilm formation and triggers the stringent response in *Staphylococcus aureus*. Sci Rep. 2017;7:40965.

10. Blazanin M, Turner P. Community context matters for bacteria-phage ecology and evolution. ISME. 2021;15:3119–28.

11. Johnke J, Baron M, de Leeuw M, Kushmaro A, Jurkevitch E, Harms H and Chatzinotas A. A Generalist Protist Predator Enables Coexistence in Multitrophic Predator-Prey Systems Containing a Phage and the Bacterial Predator *Bdellovibrio*. Front. Ecol. Evol. 2017;5:124.

12. Mumford R, Friman V. Bacterial competition and quorum-sensing signalling shape the eco-evolutionary outcomes of model in vitro phage therapy. Evol Appl. 2016;10:161–9.

13. Gómez P, Buckling A. Bacteria-phage antagonistic coevolution in soil. Science. 2011;332:106–109.

14. Middelboe M, Hagström A, Blackburn N, Sinn B, Fischer U, Borch NH, Pinhassi J, Simu K & Lorenz MG. Effects of Bacteriophages on the Population Dynamics of Four Strains of Pelagic Marine Bacteria. Microb Evol. 2001;42:395–406.

15. De Sorti L, Lourenço M, Laurent D. “I will survive”: A tale of bacteriophage-bacteria coevolution in the gut. Gut Microbes. 2019;10:92–9.

16. De Sorti L, Khanna V, Laurent D. The Gut Microbiota Facilitates Drifts in the Genetic Diversity and Infectivity of Bacterial Viruses. Cell Host Microbe. 2017;22:801–8.e3.

17. Wolfe BE, Button JE, Santarelli M, Dutton R. Cheese rind communities provide tractable systems for in situ and in vitro studies of microbial diversity. Cell. 2014;158:422–33.

18. Morin MA, Pierce EC, Dutton RJ. Changes in the genetic requirements for microbial interactions with increasing community complexity. eLife. 2018;7:e37072.

19. Pierce EC, Morin M, Little JC, Liu RB, Tannous J, Keller NP, Pogliano K, Wolfe BE, Sanchez LM, Dutton RJ. Bacterial-fungal interactions revealed by genome-wide analysis of bacterial mutant fitness. Nat Microbiol. 2021;6:87–102.

20. Adler BA, Kazakov AE, Zhong C, Liu H, Kutter E, Lui LM, Nielsen TN, Carion H, Deutschbauer AM, Mutalik VK, Arkin AP. The genetic basis of phage susceptibility, cross-resistance and host-range in *Salmonella*. Microbiol. 2021;167:001126.

21. Mutalik VK, Adler BA, Rishi HS, Piya D, Zhong C, Koskella B, Kutter EM, Calendar R, Novichkov PS, Price MN, Deutschbauer AM, Arkin AP. High-throughput mapping of the phage resistance landscape in *E. coli*. PLoS Biol. 2020;18:e3000877.

22. Kortright KE, Chan BK, Turner PE. High-throughput discovery of phage receptors using transposon insertion sequencing of bacteria. PNAS. 2020;117:18670–79.

23. van Opijnen T, Camilli A. Transposon insertion sequencing: a new tool for systems-level analysis of microorganisms. Nat Rev Microbiol. 2013;11:435–42.

24. Morin MA, Morrison AJ, Harms MJ, Dutton RJ. Higher-order interactions shape microbial interactions as microbial community complexity increases. Sci Rep. 2022;12:22640.

25. Meyer JR, Dobias DT, Weitz JS, Barrick JE, Quick RT, Lenski RE. Repeatability and contingency in the evolution of a key innovation in phage lambda. Science. 2012;335:428–32.

26. Nishimura Y, Yoshida T, Kuronishi M, Uehara H, Ogata H, Goto S. ViPTree: the viral proteomic tree server. Bioinformatics. 2017;33:2379–80.

27. Wetmore KM, Price MN, Waters RJ, Lamson JS, He J, Hoover CA, Blow MJ, Bristow J, Butland G, Arkin AP, Deutschbauer A. Rapid Quantification of Mutant Fitness in Diverse Bacteria by Sequencing Randomly Bar-Coded Transposons. mBio. 2015;6:e00306–15.

28. Eren AM, Kiefl E, Shaiber A, Veseli I, Miller SE, Schechter MS, Fink I, Pan JN, Yousef M, Fogarty EC, Trigodet F, Watson AR, Esen ÖC, Moore RM, Clayssen Q, Lee MD, Kivenson V, Graham ED, Merrill BD, Karkman A, Blankenberg D, Eppley JM, Sjödin A, Scott JJ, Vázquez-Campos X, McKay LJ, McDaniel EA, Stevens SLR, Anderson RE, Fuessel J, Fernandez-Guerra A, Maignien L, Delmont TO, Willis AD. Community-led, integrated, reproducible multi-omics with anvi’o. Nat Microbiol. 2021;6:3–6.

29. Lex A, Gehlenborg N, Strobelt H, Vuillemot R, Pfister H. UpSet: Visualization of Intersecting Sets. IEEE Trans Vis Comput Graph. 2014;20:1983–92.

30. Conway JR, Lex A, Gehlenborg N. UpSetR: an R package for the visualization of intersecting sets and their properties. Bioinformatics. 2017;33:2938–40.

31. Kanehisa M, Sato Y, Morishima K. BlastKOALA and GhostKOALA: KEGG Tools for Functional Characterization of Genome and Metagenome Sequences. J Mol Biol. 2016;428:726–31.

32. Cai R, Wang G, Le S, Wu M, Cheng M, Guo Z, Ji Y, Xi H, Zhao C, Wang X, Xue Y, Wang Z, Zhang H, Fu Y, Sun C, Feng X, Lei L, Yang Y, ur Rahman S, Liu X, Han W, Gu J. Three Capsular Polysaccharide Synthesis-Related Glucosyltransferases, GT-1, GT-2 and WcaJ, Are Associated With Virulence and Phage Sensitivity of *Klebsiella pneumoniae*. Front Microbiol. 2019;10:1189.

33. Caroff M, Novikov A. Lipopolysaccharides: structure, function and bacterial identification. OCL. 2020;27:31.

34. Letarov AV, Kulikov EE. REVIEW: Adsorption of Bacteriophages on Bacterial Cells. Biochemistry (Moscow). 2017;82:1632–58.

35. Silva JB, Storms Z, Sauvageau D. Host receptors for bacteriophage adsorption. FEMS Microbiol Lett. 2016;363:fnw002.

36. Klena JD, Ashford RS II, Schnaitman CA. Role of *Escherichia coli* K-12 *rfa* genes and the *rfp* gene of *Shigella dysenteriae* 1 in generation of lipopolysaccharide core heterogeneity and attachment of O antigen. 1992;174:7297–307.

37. Pagnout C, Sohm B, Razafitianamaharavo A, Caillet C, Offroy M, Leduc M, Gendre H, Jomini S, Beaussart A, Bauda P, Duval JFL. Pleiotropic effects of *rfa*-gene mutations on *Escherichia coli* envelope properties. 2019;9:9696.

38. Gu Z, Eils R, Schlesner M. Complex heatmaps reveal patterns and correlations in multidimensional genomic data. 2016;31:2847–2849.

39. Gu Z. Complex heatmap visualization. iMeta. 2022;1:e43.

40. Svetlov V, Belogurov GA, Shabrova E, Vassylyev DG, Artsimovitch I. Allosteric control of the RNA polymerase by the elongation factor RfaH. Nucleic Acids Res. 2007;35:5694–705.

41. Iguchi A, Iyoda S, Kikuchi T, Ogura Y, Katsura K, Ohnishi M, Hayashi T, Thomson NR. A complete view of the genetic diversity of the *Escherichia coli* O-antigen biosynthesis gene cluster. DNA Res. 2015;22:101–7.

42. Thomsen LE, Chadfield MS, Bispham J, Wallis TS, Olsen JE, Ingmer H. Reduced amounts of LPS affect both stress tolerance and virulence of *Salmonella enterica* serovar Dublin. FEMS Microbiol Lett. 2003;228:225–31.

43. Broeker NK, Barbirz S. Not a barrier but a key: How bacteriophages exploit host’s O-antigen as an essential receptor to initiate infection. Mol Microbiol. 2017:105;353–357.

44. Ebbensgaard A, Mordhorst H, Aarestrup FM, Hansen EB. The Role of Outer Membrane Proteins and Lipopolysaccharides for the Sensitivity of *Escherichia coli* to Antimicrobial Peptides. Front Microbiol. 2018;9:2153.

45. Burmeister AR, Fortier A, Roush C, Lessing AJ, Bender RG, Barahman R, Grant R, Chan BK, Turner PE. Pleiotropy complicates a trade-off between phage resistance and antibiotic resistance. PNAS. 2020;117:11207–16.

46. McGee LW, Barhoush Y, Shima R, Hennessy M. Phage-resistant mutations impact bacteria susceptibility to future phage infections and antibiotic response. Ecol Evol. 2023;13:e9712.

47. Jayaratne P, Bronner D, MacLachlan PR, Dodgson C, Kido N, Whitfield C. Cloning and analysis of duplicated *rfbM* and *rfbK* genes involved in the formation of GDP-mannose in *Escherichia coli* O9:K30 and participation of *rfb* genes in the synthesis of the group I K30 capsular polysaccharide. J Bacteriol. 1994;176:3126–39.

