## Supplemental Figures 1-3 for "O-antigen biosynthesis mediates evolutionary trade-offs within a simple community"

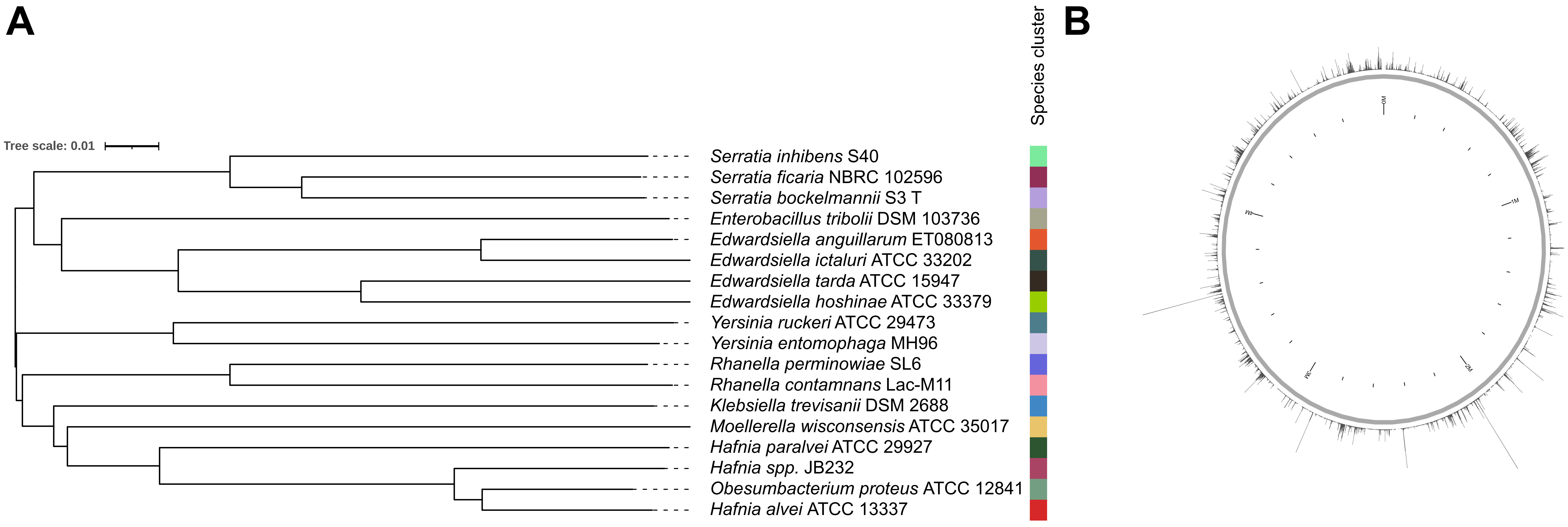


**Supplementary Figure 1. Genomic analysis and manipulation of cheese bacterial strain JB232.** (A) Genome Based Distance Phylogeny (GBDP) tree based on genome data reveals that JB232 forms a unique species cluster within the genus *Hafnia*. Phylogeny was determined using the Type (Strain) Genome Server (TYGS) and the tree was visualized using Interactive Tree of Life (iTOL). (B) Insertion map of JB232 genome library. The RB-TnSeq library was constructed in the laboratory via conjugation with *Escherichia coli* strain APA766 carrying the pKMW7 Tn5 vector library (Wetmore et al. 2015). Each vertical bar in the map represents the number of insertions within a 1000 bp region; bar height is directly proportional to the number of insertions, with approximately 1700 insertions being represented by the longest bar. Library sequencing revealed 103169 insertions in 58869 distinct locations within the JB232 genome. These insertions occurs within the central 10-90% of each represented gene. The RB-TnSeq library contains mutants in approximately 88% of the protein-coding genes, each of which has an average of 25.6 strains. The insertion map was visualized using anvi’o (Eren et al. 2021).

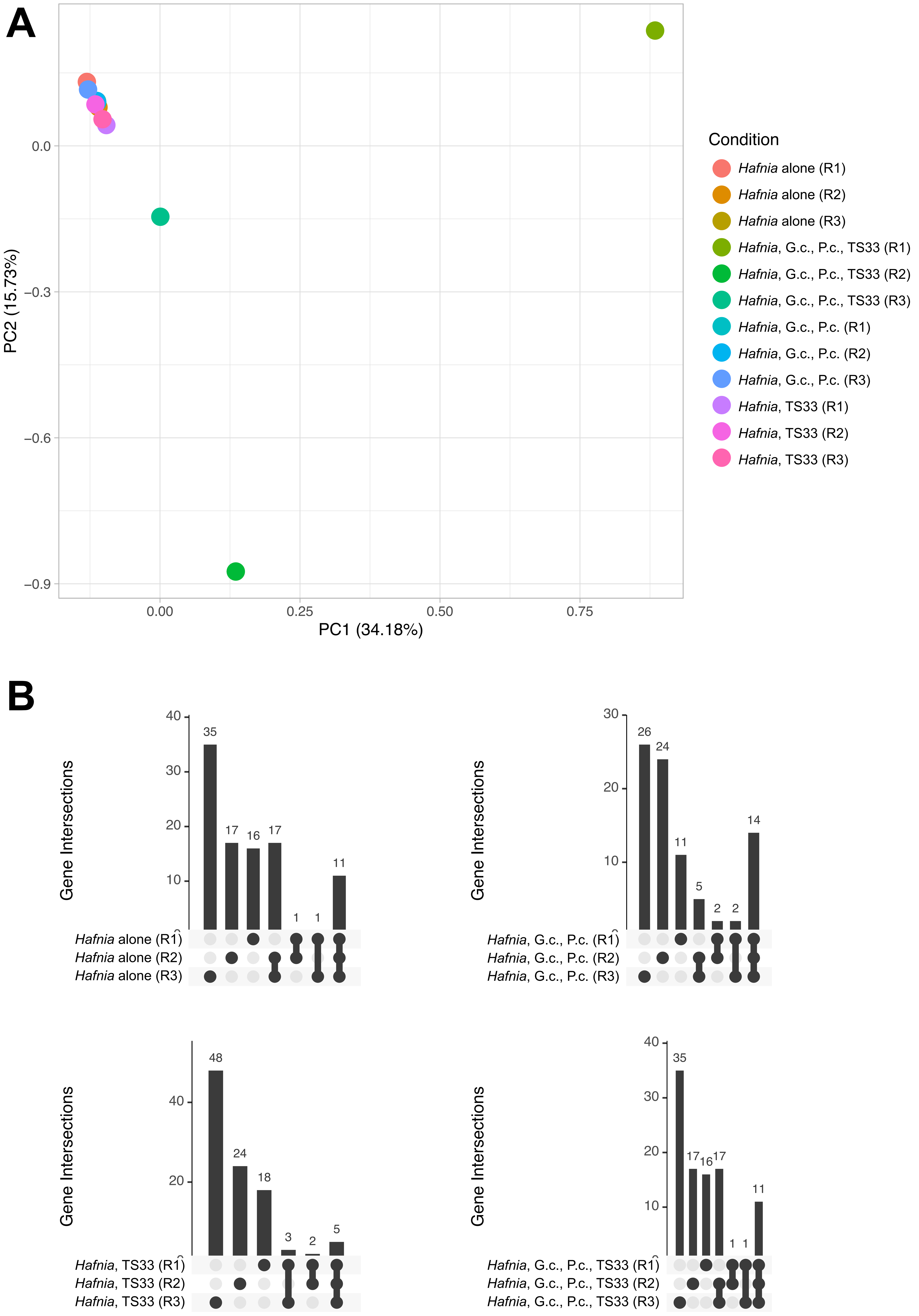


**Supplementary Figure 2**. **Comparison of significant gene fitness values for the individual replicates in each experimental condition.** (A) Principal Component Analysis was conducted on the significant gene fitness values using the prcomp() function in Rstudio. (B) We identified genes of significant fitness unique to and/or shared by the three replicates from four growth conditions using the UpSetR package in Rstudio (Lex et al. 2014, Conway et al. 2017).

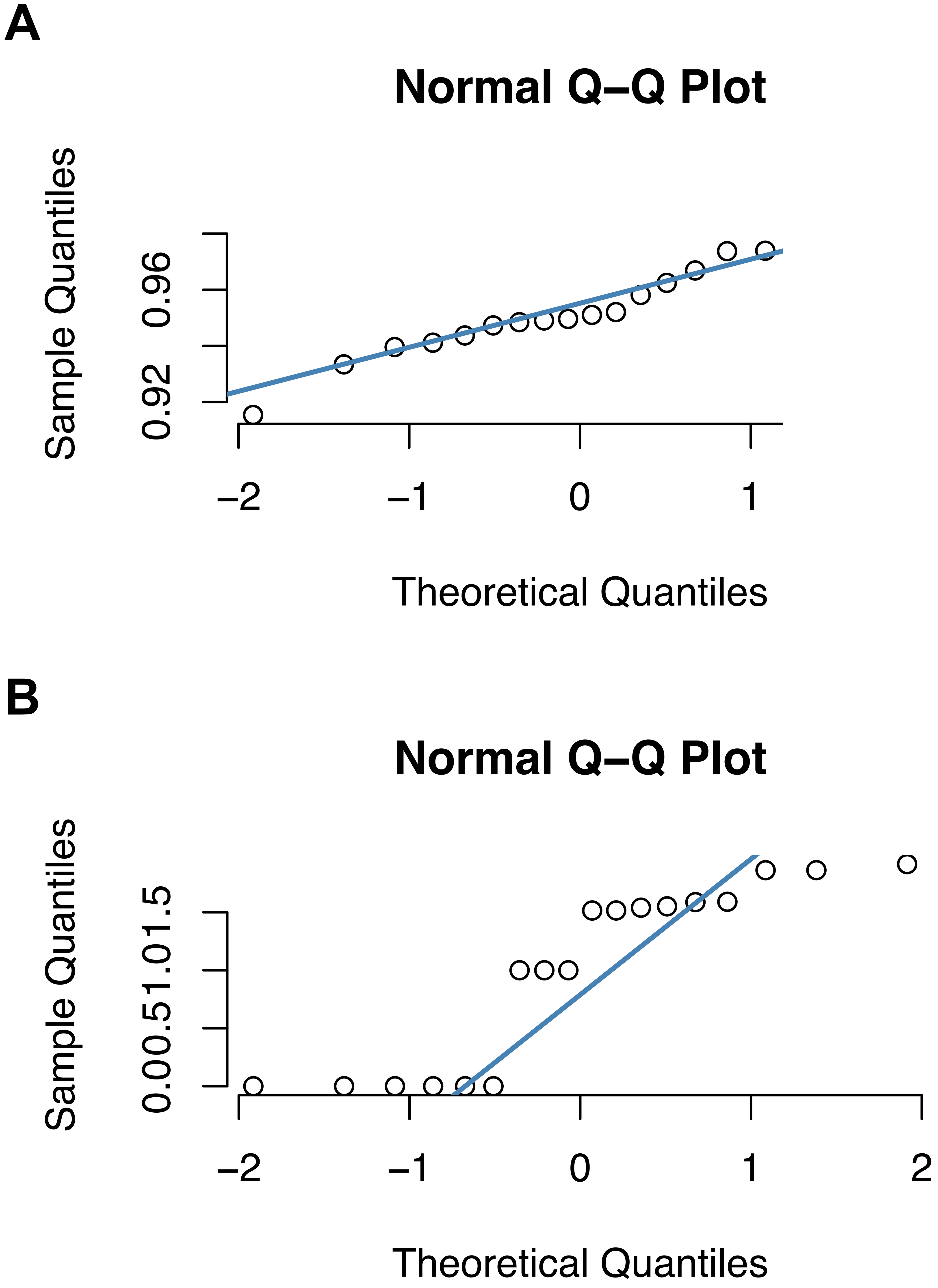


**Supplementary Figure 3**. Determining the normality of the data sets visualized in Figure 4. (A) Under all tested conditions, the number of *Hafnia* colony forming units follows a normal distribution. Shapiro-Wilk (S-W) and Pearson Chi-Squared Normality tests also produced p-values >0.05. (B) The number of plaque forming units of TS33 does not follow a normal distribution based on the shape of the quantile-quantile plot, and the p<0.05 for the S-W and Pearson Normality tests.
